# Halfpipe: a tool for analyzing metabolic labeling RNA-seq data to quantify RNA half-lives

**DOI:** 10.1101/2024.09.19.613510

**Authors:** Jason M. Müller, Elisabeth Altendorfer, Susanne Freier, Katharina Moos, Andreas Mayer, Achim Tresch

## Abstract

We introduce Halfpipe, a tool for analyzing RNA-seq data from metabolic RNA labeling experiments. Its main features are the absolute quantification of 4sU-labeling-induced T>C conversions in the data as generated by SLAM-seq, calculating the proportion of newly synthesized transcripts, and estimating subcellular RNA half-lives. Halfpipe excels at correcting critical biases caused by typically low labeling efficiency. We measure and compare the RNA metabolism in the G1 phase and during the mitosis of synchronized human cells. We find that RNA half-lives of constantly expressed RNAs are similar in mitosis and G1 phase, suggesting that RNA stability of those genes is constant throughout the cell cycle. Our estimates correlate well with literature values and with known RNA sequence features. Halfpipe is freely available at https://github.com/IMSBCompBio/Halfpipe

## 1 Introduction

The metabolism of an mRNA can be subdivided into three major processes: transcription, translation, and degradation. In contrast to prokaryotes, transcription and translation are spatially separated by the compartmentalization of a eukaryotic cell into the nucleus and cytosol. Consequently, nuclear RNA export is another step in the eukaryotic mRNA metabolism. Each step is highly regulated, ensuring a cell’s timely and flexible response to internal and external stimuli [1, 2, 3]. Impairments in mRNA metabolism have been associated with a range of human diseases including cancer and neurological diseases such as motor neuron disease [4, 5, 6]. Although of apparent interest, quantifying these metabolic processes is beset with considerable difficulties.

Transcript dynamics can be studied by metabolical RNA labeling with nucleoside analogs such as 4-thiouridine (4sU) [7, 8, 9]. When cells are exposed to 4sU, RNA polymerase II (Pol II) incorporates this analog into newly synthesized transcripts, enabling differentiation between recently synthesized (“new”) and pre-existing (“old”) transcripts. These transcript populations can be biochemically separated via biotinylation and quantified by RNA sequencing [10, 11, 12, 13]. Several studies estimated RNA synthesis and degradation rates by monitoring the fractions of these quantities, the 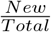 RNA ratios, over time [10, 11, 12, 13].

Biotinylation-based approaches generally suffer from the limited separation efficiency of the labeled and unlabeled RNA fraction [14], biasing downstream estimates of RNA metabolism. Recently, protocols such as SLAM-seq and Timelapse-seq have been introduced that bypass the biochemical separation [7, 15]. According to these protocols, the transcript populations are not physically separated, but 4sU is biochemically converted so that the sequencing reads contain a cytosine instead of a thymine at the 4sU incorporation site of the corresponding transcripts. These T>C conversions are sufficient to distinguish between sequencing reads derived from newly synthesized and pre-existing transcripts. Still, the data analysis is challenging due to various limitations. For instance, T>C conversions can also arise from single-nucleotide polymorphisms (SNPs) and RNA editing events [16, 17]. Also, a 4sU-labeling-induced T>C conversion could be masked by C>non-C sequencing errors. Yet, a major challenge results from the generally low 4sU incorporation rate, which is reportedly in the range of only 2% [7]. Due to this low labeling efficiency, there is a high chance that a certain fraction of the newly synthesized transcript remains unlabeled, so a naive count of T>C conversion would lead to an underestimation of the newly synthesized fraction. Such a bias can severely distort the downstream estimates of the dynamic parameters of mRNA metabolism.

We recently developed a modeling framework that overcomes many of the limitations in the analysis of metabolic RNA sequencing data [18]. Building upon this work, we present Halfpipe, a user-friendly package for quantifying transcript-specific metabolic rates on a genome-wide scale. Using data from whole-cell extracts or subcellular (e.g. nuclear and cytosolic) fractions, Halfpipe performs thorough data cleaning steps, including read mapping, quality filtering, and T>C conversion quantification. It employs an improved expectation-maximization algorithm to yield precise 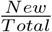 RNA ratios (Figure 1). Subsequently, a differential equation system is used to estimate rates of RNA metabolism, including synthesis, export, and degradation rates from these ratios. Halfpipe computes these rates in a post-processing phase into whole-cell or nuclear and cytosolic RNA half-lives. Finally, it applies various quality filters before returning a list of transcript and compartment-specific RNA half-lives for each measured transcript to the user. Halfpipe is freely available, along with detailed documentation, at https://github.com/IMSBCompBio/Halfpipe.

**Figure 1:**
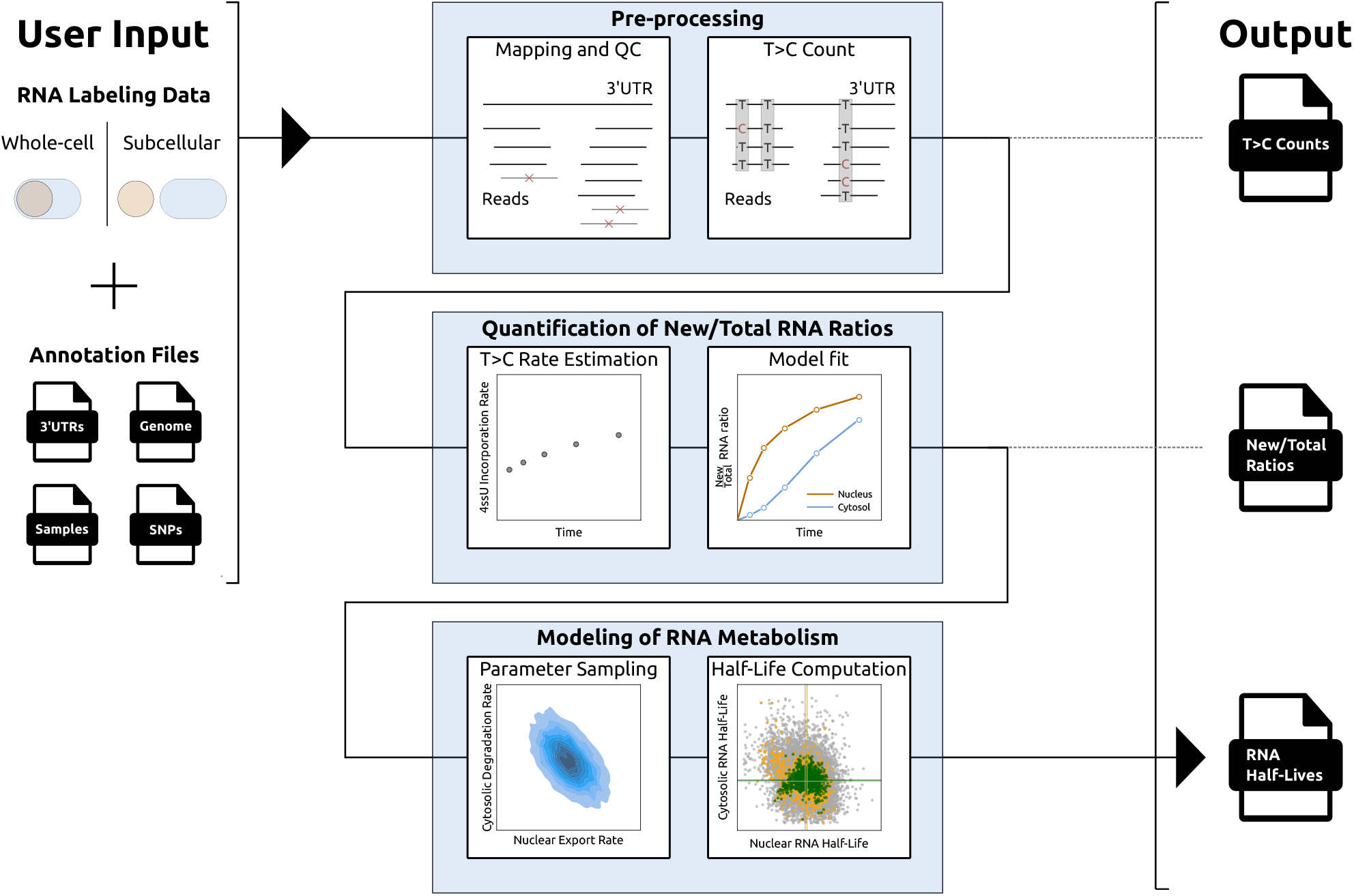
Halfpipe Workflow. The input consists of a sample sheet, a reference genome, an annotation file with genomic coordinates of 3’UTRs, and an annotation file with known SNPs. Data pre-processing includes read mapping and quality filtering to the reference genome, T>C calling under the exclusion of known and sample-specific potential SNPs and RNA editing sites. The intermediate result is a T>C conversion count table. Halfpipe estimates the 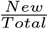 RNA ratios in each sample during the modeling phase. Finally, the parameters of a one-or two-compartment model of mRNA metabolism are fitted to these ratios using Markov Chain Monte Carlo (MCMC) sampling and optimization. This results in RNA metabolic rate estimates for each compartment and transcript and extensive quality metrics.

## 2 Methods

### 2.1 Pre-processing of RNA-seq Data

Halfpipe uses the T>C conversion aware alignment tool Nextgenmap [19] for sequencing data alignment. By default, it uses the following parameters: `-512-b--slam-seq2--max-polya4--strata-l`. Reads are filtered out if their mapping quality value is lower than 2 and their sequence identity is lower than 95%. Reads that pass these filters are assigned to known 3’UTRs according to a user-specified BED file. SNPs that could mimic 4sU-labeling-induced T>C conversions are masked based on a user-specified VCF file. Additionally, Halfpipe calls potential SNPs and RNA editing sites by calculating position-wise mismatch rates for each covered genomic position in control (i.e., no 4sU labeling) samples. If the mismatch rate is higher than 5%, the respective genomic position is masked. Subsequently, the T>C conversions are quantified from each sequencing read.

### 2.2 Quantification of New/Total RNA Ratios

For sake of generality, we present the multi-compartment model for arbitrary many compartments in the Supplements. It may serve others to expand the model to more than 2 compartments, as done by Ietswaart *et al*. (2024) and Steinbrecht *et al*. (2024) [20, 21]. Our treatment of the subject combines the presentation in Chappleboim *et al*. (2022) [22], which includes exponential cell growth, with the elegant formulation of the model in Steinbrecht *et al*. (2024) [20] (Supplements).

In standard applications, Halfpipe models the life cycle of a transcript by fitting a two-compartment model when provided with subcellular fractionation data of the nuclear and cytosolic fractions. In this model, the time course of nuclear RNA *N* and the cytosolic RNA *C* is described by the synthesis rate *µ*, the nuclear degradation rate *ν*, the nuclear export rate *τ*, and the cytosolic degradation rate *λ*, assuming a growth rate that is negligible compared to the degradation and export rates:

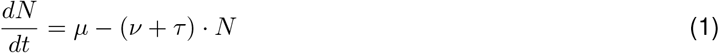

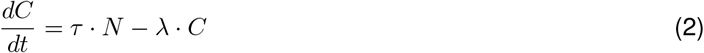

If whole-cell extract data is provided, Halfpipe applies a one-compartment model describing the time course of total RNA *R* by the synthesis rate *µ* and the degradation rate δ.

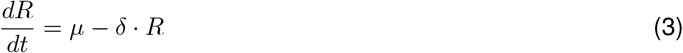

Under steady-state conditions, the two-compartment model has the closed-form solutions:

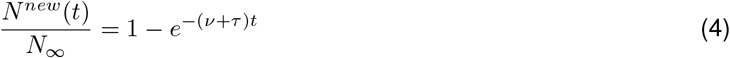

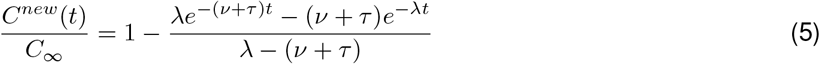

Notably, Equations (4)-(5) only depend on λ and the joint parameters ν + τ, a quantity we call nuclear vanishing rate. Correspondingly, the one-compartment model has the following solution:

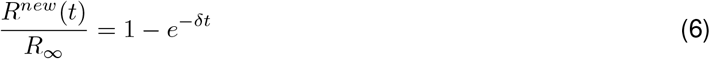

As described by Equations (4)-(6), the 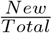 RNA ratios are the targets for Halfpipe’s model fitting procedure. However, the naive estimates of the experimentally determined ratios are biased due to the low 4sU incorporation rate of roughly 2% [7]. Similar to Jürges *et al*. (2018) [23], Halfpipe models the actual 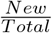 RNA ratios using a binomial mixture model

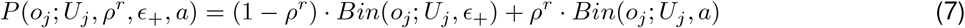

where *o*_*j*_ is the observed number of T>C conversions and *U*_*j*_ is the number of potential conversion sites in a read *j* assigned to a particular 3’UTR *r* ∈ ℛ. *ρ*^*r*^ is the 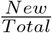 RNA ratio of region *r*, and *ϵ*_+_ is the T>C false-positive sequencing error. *a* denotes a corrected labeling efficiency, which, besides the actual labeling efficiency *ℓ*, takes the T>C false-positive sequencing error *ϵ*_+_ and the C>X false-negative sequencing error *ϵ*_−_ (error rates estimated from control samples as described in Müller *et al*. (2024) [18]) into account:

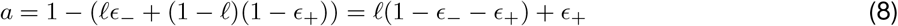

Halfpipe estimates the unknown parameter set Θ = (*ℓ, ρ*^*r*^, *r* ∈ ℛ) with an Expectation-Maximization (EM) algorithm (Supplements). Subsequently, *ℓ* is plugged into Equation (8), which in turn is plugged into the binomial mixture model described in Equation (7). Then, Halfpipe performs a last round of optimization (L-BFGS-B method [24]) yielding a final 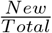 RNA ratio estimate *ρ*^*r*^ for each 3’UTR *r*.

### 2.3 Modeling of RNA Metabolism

Having determined the 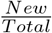 RNA ratios, Halfpipe applies an arcsin-transformation [25] to these quantities as a variance-stabilization step before model fit (Supplements). Then, either a two-or one-dimensional Metropolis-Algorithm (MCMC method) is used to fit Equation (4)-(5) (two-compartment model) or Equation (6) (one-compartment model), respectively. A seed for the Markov chain is determined by global optimization (Differential Evolution algorithm [26]). We sample 50,000 parameter combinations and use their central 95%-interval as confidence intervals. Using median values of the MCMC samples, the final parameter set Θ = (ν + *τ, λ*) (two-compartment model) or *δ* is determined using local optimization (two-compartment model: Nelder-Mead algorithm [27]; one-compartment model: Brent algorithm [28]).

The reciprocal values of the metabolic RNA rate estimates, 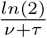 and 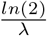 are the nuclear and cytosolic RNA half-lives (two-compartment model) of a respective transcript. Correspondingly, 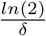 describes the whole-cell RNA half-life (one-compartment model). Further, Halfpipe classifies an estimate as highly reliable if it fulfills the following criteria: (1) The corresponding 3’UTR has an average read count of at least 30 along the time series experiment. (2) The expression level is constant (steady-state assumption). That means the average slope of a linear model fit to the expression levels over time does not exceed a value of 0.0025 (or does not fall below a value of -0.0025) (3) The deviation between the parameter estimate and the upper and lower credibility limits by the parameter is ≤ 0.3. (4) The model fit is reasonable (*R*^2^ ≥ 0.4).

### 2.4 SLAM-seq Cell Cycle Data

HeLa-S3 cells were cultured at 37°C in DMEM supplemented with 10% FBS Superior (Sigma, S0615) and 1X penicillin-streptomycin (Gibco, 15070063). A double thymidine block was used for cell cycle synchronisation at the G1/S boundary. Briefly, Hela-S3 cells were blocked with 2.5 mM thymidine for 18 hours. The cells were released from the block by two washes with pre-warmed medium, followed by incubation for 9 hours in fresh medium supplemented with 24 µM 2-deoxycytidine. The second G1/S block was again performed by adding 2.5 mM thymidine for 18 hours. Finally, cells were released by two washes with pre-warmed media and incubated for 6 hours (late G2) in DMEM containing 24 µM 2-deoxycytidine before labelling with 500 µM 4sU (Glentham Life Sciences, GN6085) for 0, 15, 45, 75, 120 and 180 minutes. At each time point, 3 million cells were fractionated to obtain nuclear and cytosolic fractions based on Mayer *et al*. [29] with the following modifications: Nuclear pellets were washed with nuclear wash buffer (PBS with 1X cOmplete Protease Inhibitor (Roche, 11873580001)) and resuspended in 1400 µl TRIzol (Invitrogen, 15596026). Similarly, a time series in the G1 cell cycle phase (12 hours release time after double thymidine blocking) was measured at 0, 15, 45, 75, 145, 180, 240 and 300 minutes. Prior to phenol extraction of RNA, 1.9 µl of 1:100 diluted ERCC ExFold RNA Spike-In Mixes (Invitrogen, 4456739) per 1 million cells was added to each sample. The 4sU RNA was alkylated according to Herzog *et al*. (2017) [7], and poly-A enriched SLAM-seq libraries were generated using the 3’-QuantSeq-REV protocol (Lexogen) according to the manufacturer’s protocol. The final libraries were sequenced on an Illumina NovaSeq6000 sequencer at 150 nucleotide read lengths in paired-end mode. Each time series was used as input for Halfpipe together with a GRCh38 reference genome file, a 3’UTR annotation file from UCSC [30] (overlapping regions were merged), and a VCF with SNP positions from NCBI dbSNP [31]. Only the first read of each mapped read pair was taken into account for subsequent analyses to avoid double counting of T>C conversions from possibly overlapping read pairs.

### 2.5 Simulations

For Figure 2A, 1,000 3’UTRs were simulated with a fixed coverage *T* of 10, 30, 100, 200, 500, and 1,000 reads per scenario. Each read was simulated to contain 10, 20, 30, 40, and 50 potential 4sU labeling sites *U*. The 4su labeling efficiency *ℓ* was set to a literature value of 2% [7]. The false-positive T>C error *ϵ*_+_ and the false-negative C>x error *ϵ*_−_ were set to a realistic sequencing error of 0.1% [32]. Each region was assigned a uniformly sampled 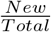 RNA ratio *ρ*. Then, the reads were classified into two groups: the newly synthesized fraction *N* ∼ *Bin*(*T, ρ*) and, subsequently, the pre-existing fraction *O* = *T* − *N*. Then, T>C conversions in new reads were sampled from a binomial distribution *Bin*(*U, a*) (*a* is the corrected labeling efficiency as described by Equation (8)) and in old reads from *Bin*(*U, ϵ*_+_). These T>C conversion counts were subjected to Halfpipe’s EM algorithm (Supplements) to estimate the labeling efficiency *ℓ*.

**Figure 2:**
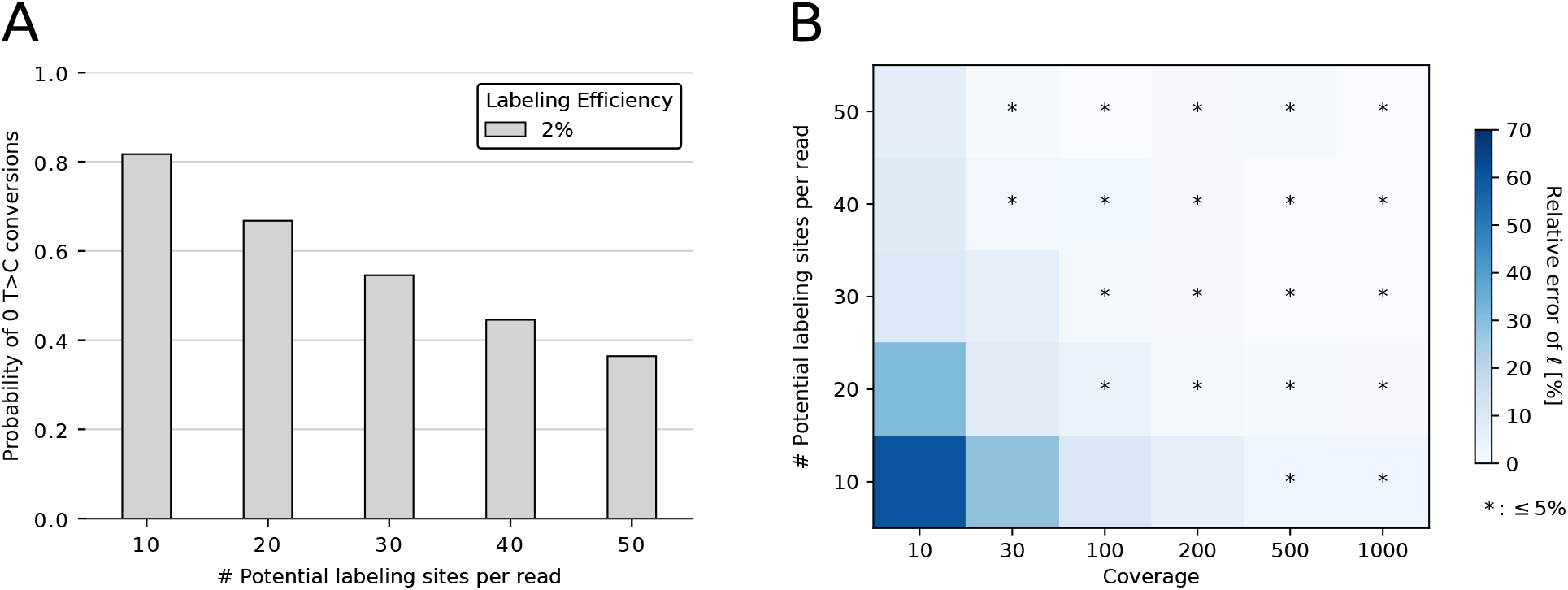
**(A)** Probability of observing no T>C conversions in a sequencing read (y-axis) with increasing numbers of potential 4sU incorporation sites (x-axis), assuming site-independent labeling and a labeling efficiency of 2%. **(B)** Heatmap showing the relative error of the labeling efficiency estimates by Halfpipe. Darker colors indicate higher relative error rates (color bar).

To study the impact of deviations in the 4sU incorporation rate (Figure 3A), 3’UTRs were simulated from annotated transcripts of the GRCh38 human reference genome (annotation file downloaded from the UCSC Table Browser [30]). 1,000 3’UTRs, each with a coverage *T* ^*r*^ ranging between 0-1000, were generated. Here, reads were sampled from a Beta-Binomial *BetaBin*(*n* = 1000, *α* = 90, *β* = 10) to obtain an average of 100 reads per region *r*. The length of each read was set to 100 nt. Each read was sampled from within the last 250 nt of annotated transcripts. Consequently, the number of potential labeling sites could vary between reads from the same region *r*. Next, a 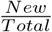 RNA ratio *ρ*^*r*^ ∈ {1%, 5%, 25%.50%, 75%, 95%, 100%}. The number of new reads *N* ^*r*^ was then sampled from a binomial *Bin*(*T* ^*r*^, *ρ*_*r*_) so that the number of old reads *O*^*r*^ was *O*^*r*^ = *T* ^*r*^ −*N* ^*r*^. Again, the labeling efficiency *ℓ* was set to 2%, the false-positive T>C error *ϵ*_+_ to 0.1%, and the false-negative C>X error *ϵ*_−_ both to 0.1%. Then, for each read *j* ∈ *T* ^*r*^, the number of potential 4sU incorporation sites 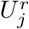 was counted. Subsequently, T>C conversions were sampled from 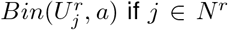 (new fraction) or 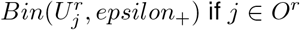 (pre-existing fraction). Here, *a* denotes the corrected labeling efficiency (Equation (8)). The T>C conversion counts were then used to estimate *ρ*. However, the labeling efficiency was manually varied between 1-3% with a step size of 0.01%. At each step, *a* was calculated (Equation (8)), which was plugged into the binomial mixture model (Equation (7)) to yield an 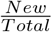 RNA ratio estimate *ρ*^*r*^ for each region *r*.

**Figure 3:**
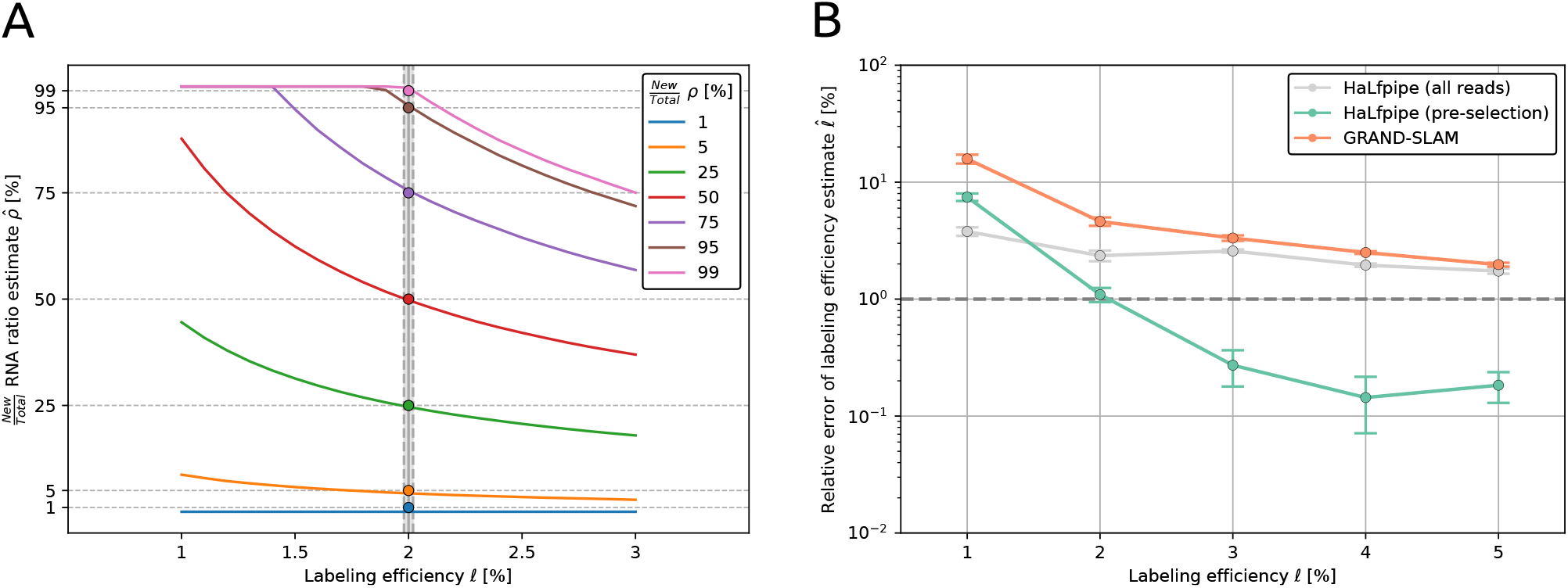
**(A)** Sensitivity of the 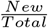 RNA ratio estimates as a function of the labeling efficiency. The data was generated with an actual labeling efficiency of 2% (grey vertical line) for a range of 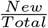 NA ratios (colored dots). The 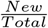 RNA ratios were reestimated while the labeling efficiency was intentionally misspecified as indicated on the x-axis. **(B)** Benchmarking of Halfpipe against GRAND-SLAM’s [23] EM algorithm (orange line) performance regarding 4sU incorporation rate estimation. Halfpipe was run either with all reads (gray line) or with a pre-selected fraction whose reads contain at least 30 potential 4sU incorporation sites (green line). Points represent the means of 5 replicates in each labeling efficiency scenario, and error bars show the standard error of the mean. The horizontal dotted line marks a relative error of 1% in the labeling efficiency estimate deemed acceptable.

The simulation for the performance assessment of Halfpipe (Figure 3B) was similar to the study of the impact of deviations in the 4sU incorporation rate (Figure 3A), but 15,000 3’UTRs were simulated from annotated transcripts of GRCh38 (annotation file downloaded from the UCSC Table Browser [30]). *ϵ*_+_ and *ϵ*_−_ were set to 0.1%, 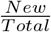 RNA ratios *ρ*_*r*_ ranging between 1-99% was sampled uniformly for each region *r*, and reads sampled from a Beta-Binomial *BetaBin*(*n* = 1000, *α* = 90, *β* = 10). For each region *r*, reads originating from new transcripts *N* ^*r*^ were sampled from a binomial *Bin*(*T* ^*r*^, *ρ*_*r*_) so that the number of old reads *O*^*r*^ was *O*^*r*^ = *T* ^*r*^ − *N* ^*r*^. The number of potential 4sU incorporation sites 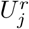 was determined for each read *j*. Subsequently, T>C conversions were sampled by 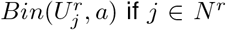 (new fraction) or 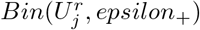 (pre-existing fraction). Here, *a* denotes the corrected labeling efficiency, which accounts for the labeling efficiency *ℓ* ∈ {1%, 2%, 3%, 4%, 5%} (Equation (8)). Halfpipe’s EM algorithm was then used with and without read pre-selection to estimate *ℓ*. The corresponding methodology of the tool GRAND-SLAM was used for comparison in terms of the accuracy of the labeling efficiency estimation. To that end, GRAND-SLAM’s EM algorithm was implemented as described in Jürges *et al*. (2018) [23]. A naive labeling efficiency of 100% was used as a start value for this EM.

## 3 Results

### 3.1 Functionality of Halfpipe

An RNA-seq experiment with metabolic RNA labeling aims to answer three questions: Which positions in the RNA were labeled by the uridine analog? What is the proportion of a gene’s labeled (i.e., recently synthesized) transcripts within the labeling period? What can be learned about the dynamics of RNA metabolism, i.e., what are the (compartment-specific) half-lives of a transcript? Halfpipe takes 4sU-seq, SLAM-seq, or TimeLapse-seq data as primary input and addresses the above questions in one automated workflow (Figure 1).

Halfpipe requires four input files that the user must provide: a sample sheet, a reference genome, and annotation files for 3’UTR regions as well as single-nucleotide polymorphisms (SNPs). Optionally, the user may specify settings such as the number of CPU cores for parallel processing. Halfpipe starts with the sequence alignment using NextGenMap [19]. The mapped reads are further quality filtered and assigned to the 3’UTRs according to the user-provided annotation file. Before counting 4sU-labeling-induced T>C conversions, SNPs that could mimic T>C conversions are masked. Additionally, potentially unannotated SNPs and RNA editing sites are called and masked using control samples as described in Müller *et al*. (2024) [18]. Halfpipe provides the resulting T>C conversion counts in a separate output file to the user. Next, Halfpipe estimates sample-specific 4sU incorporation rates -a crucial step for subsequent quantification of the newly synthesized RNA fraction. As the 4sU incorporation rate is typically only in the range of 2% [7], Halfpipe fits a statistical model to account for biases resulting from low labeling efficiency (Methods, Supplements and section below) and returns the estimated 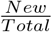 RNA ratios as a separate output. The ratios are used as targets for fitting either a one-or two-compartment model of RNA metabolism using Markov chain Monte Carlo (MCMC) sampling and optimization (Methods). This procedure returns RNA metabolic parameters such as nuclear export and degradation rates for every measured 3’UTR, respectively, their half-lives. Lastly, the tool applies stringent quality measures regarding model fit, error bounds, and constant expression (Methods). As a result, the user is provided with a summary file listing each RNA half-life and its associated quality measures.

### 3.2 Determining the Proportion of Newly Synthesized Transcripts

A major challenge in RNA labeling experiments is the generally low 4sU incorporation rate of roughly 2% [7]. As a consequence, newly synthesized transcripts might contain a low number of T>C conversions or, in the worst case, none at all. A naïve counting of T>C conversions would lead to a severe underestimation of the proportion of newly synthesized RNAs. Previously, we and others used a two-binomial mixture to model the number of conversion counts in new and pre-existing RNA [18, 21, 23]. The fitting of the model parameters by an EM algorithm was beset with considerable difficulties due to the excessive number of non-labeled yet newly synthesized transcripts [23] (Methods). However, we noticed that the chance of observing no T>C conversions decreases with the number of potential labeling sites in a sequencing read (Figure 2A). Therefore, we circumvent the underestimation of the newly synthesized fraction by selecting a subset of reads for the EM algorithm. This selection only contains reads whose numbers of potential 4sU incorporation sites exceed a certain threshold. To determine a suitable threshold, we simulated 1,000 3’UTRs spanning a broad range of 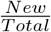 RNA ratios and coverages (Methods) at a labeling efficiency of 2%. We then assessed Halfpipe’s capability to estimate this labeling efficiency as a function of the number of potential 4sU incorporation sites and coverage (Figure 2B).

We found that a coverage of at least 30 reads per 3’UTR with 40 4sU incorporation sites per read leads to less than 5% relative error in the labeling efficiency estimate. While increasing coverage and read length is favorable regarding accuracy, sequencing costs would inevitably increase. Considering typical read lengths ranging between 50-150 nt, the number of potential 4sU incorporation sites per read will, on average, be less than 40. As a practical guide, we therefore recommend that users apply a threshold of at least 30 4sU incorporation sites and a coverage of at least 100 reads per 3’UTR for reliable parameter estimates. This represents a compromise between estimation accuracy and experimental resources.

### 3.3 Performance Assessment

The principal value of our algorithm is the accurate estimation of the labeling efficiency. First, we show by simulations that estimation of the labeling efficiency is pivotal to all subsequent analyses. We simulated SLAM-seq / 4sU-RNA sequencing data with a known, fixed labeling efficiency of 2% from 1,000 3’UTRs of the GRCh38 human genome (Methods). Using this data, we show that even slight deviations in the labeling efficiency estimates have a substantial impact on the accuracy of the 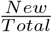 RNA ratios (Figure 3A). Generally, the 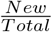 RNA ratios are noticeably underestimated if the labeling efficiency is overestimated and vice versa.

Second, scaling up the simulation to 15,000 transcripts, we benchmarked the performance of Halfpipe against its EM algorithm without pre-selection of reads with sufficiently many 4sU incorporation sites (Methods). We also included the EM algorithm implemented in GRAND-SLAM [23] into the comparison (Figure 3B). Halfpipe achieves highly accurate estimates for labeling efficiencies above 2% and outperforms GRAND-SLAM’s EM algorithm by margins across the whole range of realistic labeling efficiencies. Notably, all algorithms yield insufficient accuracy when the labeling efficiency drops below 2%. Assuming a labeling efficiency of 1%, Halfpipe performs even worse than its counterpart without read pre-selection. We conclude that – irrespective of the statistical method employed -a labeling rate of at least 2% is required for reliable quantification of 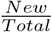 RNA ratios. Fortunately, most published studies have achieved this labeling efficiency [7, 20, 21]. On the other hand, if the estimated labelling efficiency is less than 2%, one must be aware that the data may not be sufficient for a quantitative analysis of RNA metabolism.

### 3.4 Halfpipe quantifies compartment-specific RNA half-lives

Next, we aimed to quantify the RNA metabolism throughout the cell cycle. To that end, we performed two SLAM-seq time series with HeLa-S3 cells, which were synchronized to be either in the Mitosis or G1 phase of the cell cycle (Methods). We used Halfpipe to process this data and obtain compartment-specific RNA half-lives for both cell cycle phases (Figure 4A).

**Figure 4:**
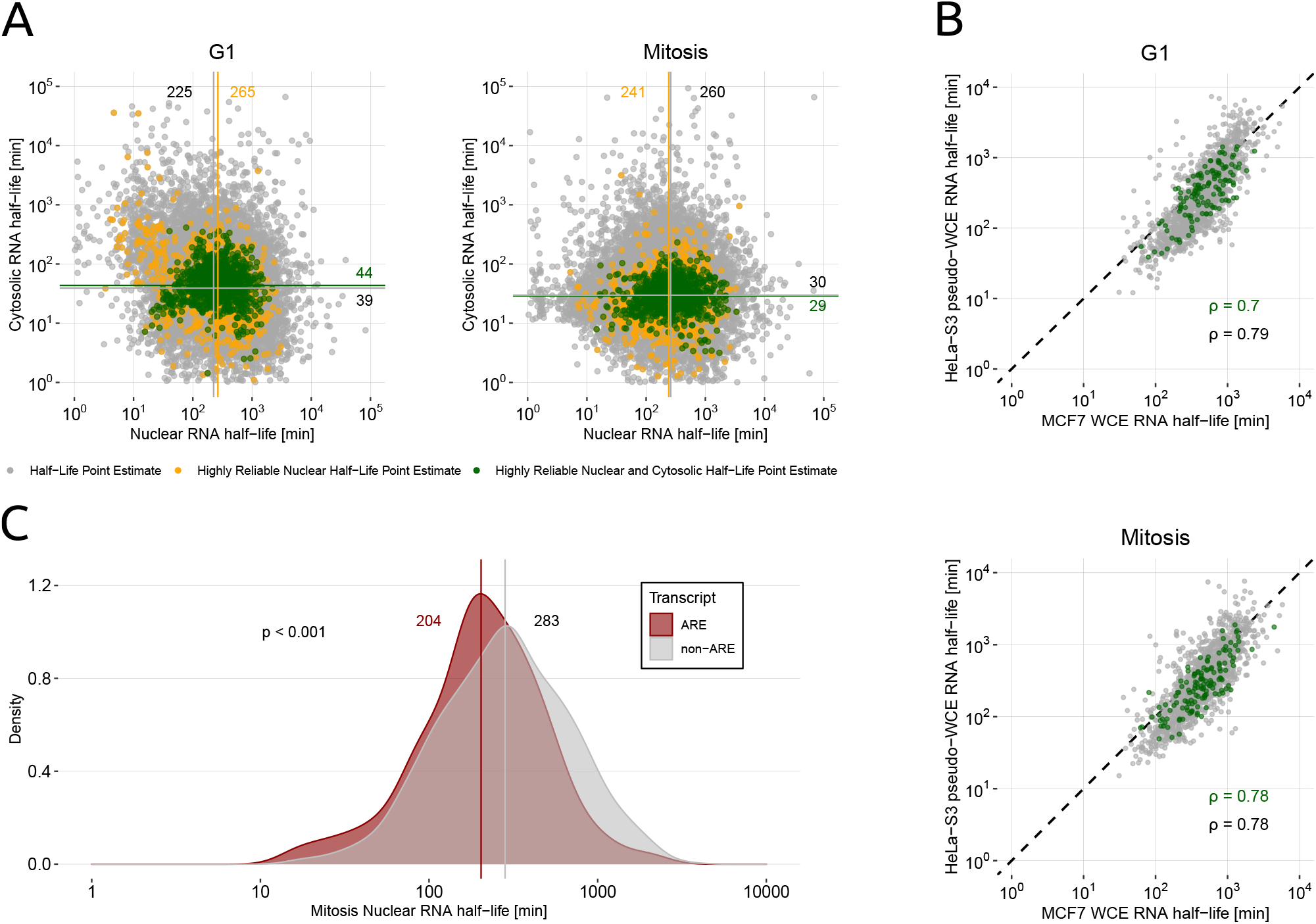
**(A)** Nuclear versus cytosolic RNA half-life measured from synchronized HeLa-S3 in G1 (left) and Mitosis (right). Gray dots reflect point estimates, while orange dots represent highly reliable nuclear and green dots represent highly reliable nuclear and cytosolic RNA half-life estimates. The horizontal and vertical lines represent the median RNA half-lives of the transcripts in these categories. **(B)** Comparison of cell cycle-specific HeLa-S3 RNA half-lives with whole-cell RNA half-lives from HEK293 cells (Schueler *et al*. (2014) [33]). Nuclear and cytosolic RNA half-lives obtained by Halfpipe were added up to pseudo-whole-cell RNA half-lives for comparison. Gray points depict point estimates, whereas green points represent highly reliable nuclear and cytosolic RNA half-lives. **(C)** Comparison of mitotic RNA half-lives of transcripts annotated to be enriched in AU-elements (red) or not (gray) according to the ARED-plus database [34]. A Wilcoxon rank-sum test was applied to assess significant differences.

We found that the average nuclear RNA half-life is 265 minutes in G1 and 241 minutes in Mitosis, considering highly reliable RNA half-life estimates. In the cytosol, it is 44 minutes in G1 and 29 minutes in Mitosis (Figure 4A). Overall, we find a high agreement between G1 and mitotic RNA half-lives of the reliable estimates (Supplemental Figure S1): the nuclear RNA half-lives’ Spearman correlation is *ρ* = 0.91 and the cytosolic one *ρ* = 0.76. These numbers suggest that the RNA metabolism of most transcripts is constant between the G1 phase and Mitosis, while also pointing towards individual transcripts deviating from this rule (Supplemental Figure S1).

We validated the half-life estimates obtained by Halfpipe by previously published half-lives obtained from whole-cell extracts of HEK293 cells [33]. The sums of our reliable nuclear and cytosolic RNA half-lives highly correlate with the whole-cell half-lives from the literature (Figure 4B; G1: *ρ* = 0.7, Mitosis: *ρ* = 0.78). Further, we verified that Halfpipe’s estimates reflect the effect of known sequential transcript features on RNA half-lives. AU-rich element (ARE) transcripts undergo increased degradation [35]. Consequently, the RNA half-lives of these transcripts should be shorter than those of non-ARE transcripts. We classified our transcripts into ARE and non-ARE RNAs based on annotation by the ARED-plus database [34] (Figure 4C). Indeed, we found significant differences (*p <* 0.001; Wilcoxon rank-sum test) between the RNA half-lives ARE and non-ARE transcripts (average ARE half-life: 204 min; non-ARE: 283 min), confirming our hypothesis.

## 4 Discussion

In this paper, we present Halfpipe, a software for quantifying RNA half-lives from metabolic labeling sequencing data. By combining data pre-processing, modeling, and post-processing, Halfpipe is easy to use. We have shown that its methodology corrects common biases in metabolic labeling sequencing data. Halfpipe’s EM algorithm provides more accurate estimates of 4sU incorporation than GRAND-SLAM’s corresponding EM methodology. The gain in performance can be attributed to improved pre-processing (filtering of SNPs) and an improved modeling of sequencing errors. The most relevant improvement was the pre-selection of reads that enter Halfpipe’s EM algorithm. While GRAND-SLAM selects reads with a high number of observed T>C conversions, we select reads with a high number of potential T>C conversions. Consequently, our pre-selection is unbiased and does not enrich for reads with high numbers of T>C conversions -which could be due to technical artifacts. Especially for short labeling pulses (resulting in a relatively low labeling efficiency and a low abundance of recently synthesized RNA), GRAND-SLAM picks only a small number of reads as input to its EM, affecting its robustness [23].

We have demonstrated that accurate labeling efficiency estimation is essential for accurate RNA half-life estimation. By applying Halfpipe to SLAM-seq data, we found that the mRNA metabolism of constantly expressed genes is largely unchanged throughout the cell cycle. Furthermore, we validated Halfpipe’s RNA half-life estimates against literature values and tested whether they reflect RNA sequence features that affect transcript stability.

We anticipate that the estimation of labeling efficiency could be further improved by replacing the underlying binomial model with a beta-binomial model in the EM algorithm. This assumption is motivated by the possibility that 4sU incorporation rates may not be constant along a transcript because of variations in Pol II elongation rate or Pol II accessibility to 4sU. A beta-binomial distribution may better capture these variations and account for the over-dispersion observed in the binomial model. Another major challenge in RNA metabolic analysis is the comparison of different transcript isoforms of a gene. For example, it would be possible to quantify the effect of nonsense-mediated decay on isoform-specific gene expression. The modular architecture of Halfpipe allows for the incorporation of an isoform-aware mapping tool. Given recent advances in sequencing technologies, measuring isoform-specific RNA half-lives is a promising venue.

In this paper we have focused on studying the nucleoplasmic and cytosolic fractions of a eukaryotic cell. However, Halfpipe can also be applied to other fractions. For instance, it could be applied to the chromatin-associated and nucleoplasmic fractions to study the dynamics of nascent RNA, as in Ietswaart *et al*. (2024) [21]. Furthermore, Steinbrecht *et al*. (2024) analyzed the nucleoplasmic, cytosolic and membrane-bound fractions in a recent preprint [20], indicating the necessity of future multi-compartment modeling. The modular architecture of Halfpipe allows for easy extension to multi-compartment models.

We are confident that Halfpipe, with its unique features and versatility, will significantly enhance the study of RNA metabolism. Its ability to correct for common biases in metabolic labeling sequencing data, provide accurate estimates of 4sU incorporation, and its potential for multi-compartment modeling make it a valuable tool for the RNA research community.

## 6 Acknowledgements

The authors thank Thanina Chabane, Zahra Nasrollah, Till Baar, Niklas Kleinenkuhnen, and Mohammad Hussainy for valuable discussions and comments on the manuscript. We thank the Sequencing facility of the MPI for Molecular Genetics for sequencing. We furthermore thank the Regional Computing Center of the University of Cologne (RRZK) for providing computing time on the DFG-funded (Funding number: INST 216/512/1FUGG) High Performance Computing (HPC) system CHEOPS as well as support. This work is based on Jason M. Müller’s PhD thesis [36].

## 7 Author Contributions

JMM, KM, and AT developed the model. JMM and KM implemented the model. JMM performed the computational data analysis. EA and SF designed and performed the SLAM-seq experiments under supervision by AM. AT supervised the computational work. JMM and AT wrote the manuscript with input from all authors.

