## Supplements for "Halfpipe: a tool for analyzing metabolic labeling RNA-seq data to quantify RNA half-lives"

### General Multi-Compartment Model

Suppose we have $n$ compartments, and let $R_i = R_i(t)$ the RNA abundance in compartment $i$, $i = 1, ..., n$. Let the total biomass $M = M(t)$ in the experiment grow exponentially at known rate $\alpha \geq 0$. Let $\dot{R}_i$ be the time derivative of $R_i$.Our multi-compartment model is defined by the ODE system

$$\begin{aligned} \dot{M} &= \alpha M \\ \dot{R}_1 &= \mu M - \lambda_1 R_1 \\ \dot{R}_i &= \lambda_{i-1} R_{i-1} - \lambda_i R_i \qquad i = 2, ..., n \end{aligned} \tag{9}$$

Letting $r_i = R_i/M$, we obtain

$$\begin{aligned} \dot{r}_1 &= \left(\frac{R_1}{M}\right)^{\cdot} = \frac{\dot{R}_1 M - R_1 \dot{M}}{M^2} \\ &= \frac{\mu M^2 - \lambda_1 R_1 M - \alpha R_1 M}{M^2} \\ &= \mu - (\lambda_1 + \alpha) \cdot r_1 \end{aligned} \tag{10}$$

and similarly

$$\dot{r}_i = \lambda_{i-1} r_{i-1} - (\lambda_i + \alpha) \cdot r_i \qquad i = 2, ..., n \tag{11}$$

Letting $r = (r_1, ..., r_n)^T$ and

$$A = \begin{pmatrix} \lambda_1 + \alpha & 0 & & & \\ -\lambda_1 & \lambda_2 + \alpha & 0 & & \\ 0 & -\lambda_2 & \ddots & & \\ & 0 & \ddots & \ddots & 0 \\ & & 0 & -\lambda_{n-1} & \lambda_n + \alpha \end{pmatrix} \tag{12}$$

we obtain the ODE

$$\dot{r} = -Ar + \mu e_1 \tag{13}$$

where $e_1 = (1, 0, ..., 0)^T \in R^n$. The corresponding homogenous ODE $\dot{r} = Ar$ has the solutions $r(t) = \exp(-tA) \cdot r^i$, $r^i \in R^n$, where the matrix exponential function is defined by the power series $\exp(X) = \sum_{i=0}^{\infty} \frac{X^i}{i!}$. For specific evaluation times $t = t_1, ..., t_T$ in time series measurements, $\exp(-t_j A)$ can be calculated easily.

One inhomogenous solution is the steady-state solution ($\dot{r} = 0$), $r^* = \mu A^{-1} e_1$ (we assume that there is a non-zero transport between each of the consecutive compartments and a non-zero decay in the last compartment, i.e., $\lambda_i > 0$, $i = 1, ..., n$. This implies that $A$ is invertible as the determinant of $A$ equals $\prod_{i=1}^{n}(\lambda_1 +$

$\alpha) > 0$). The general solution to (13) is therefore

$$r(t) = r^* + \exp(-tA) \cdot r^i \tag{14}$$

Re-writing Equation (14) in terms of $r^0 = r(0) = r^* + r^i$ ,we obtain

$$r(t) \quad = \quad r^* + \exp(-tA) \cdot (r^0 - r^*) \tag{15}$$

For newly synthesized RNA, we have $r^0 = 0$, and the relative abundance of newly synthesized RNA vs. total RNA becomes, for $i = 1, ..., n$,

$$q_i(t; \lambda, \alpha) := r_i/r_i^* = \frac{r_i^* - (\exp(-tA) \cdot r^*)_i}{r_i^*} = 1 - (\exp(-tA) \cdot r^*)_i/(r^*)_i \tag{16}$$

$$= \quad 1 - (\exp(-tA) \cdot A^{-1}e_1)_i/(A^{-1}e_1)_i$$

(note that $q_i$ does not depend on the parameter $\mu$). Given time series measurements at $t = t_1, ..., t_T$, the parameter $\lambda$ is tuned such that the values $q_i(t_j; \lambda, \alpha)$, $i = 1, ..., n$, $j = 1, ..., T$ fit to the data.

Specialising to the two-compartment model ($n = 2$), additionally assuming $\lambda_1 \neq \lambda_2$), Equation (16) can be simplified further by elemantary calculations

$$A^{-1} = \frac{1}{(\lambda_1 + \alpha)(\lambda_2 + \alpha)} \begin{pmatrix} \lambda_2 + \alpha & 0 \\ \lambda_1 & \lambda_1 + \alpha \end{pmatrix} \tag{17}$$

$$\exp(-tA) = \begin{pmatrix} e^{-t(\lambda_1+\alpha)} & 0 \\ \frac{\lambda_1}{\lambda_2 - \lambda_1} \left(e^{-t(\lambda_1+\alpha)} - e^{-t(\lambda_2+\alpha)}\right) & e^{-t(\lambda_2+\alpha)} \end{pmatrix} \tag{18}$$

Plugging these into Equation (16) leads to

$$\begin{aligned} q_1(t; \lambda, \alpha) &= 1 - e^{-t(\lambda_1+\alpha)} \\ q_2(t; \lambda, \alpha) &= \frac{1}{\lambda_2 - \lambda_1} \left((\lambda_2 + \alpha)e^{-t(\lambda_1+\alpha)} - (\lambda_1 + \alpha)e^{-t(\lambda_2+\alpha)}\right) \end{aligned}$$

### Expectation Maximization Algorithm to determine the 4sU incorporation rate

Let $R$ denote the set of all 3'UTRs that are expressed across a 4sU-RNA-seq/SLAM-seq/TimeLapse-seq time-series experiment. For $r \in \mathcal{R}$, let $J^r$ be the read count of region $r$. Given a read $j = 1, ..., J^r$, let $U_j^r$ be its number of potential T>C conversion sites and let $o_j^r \in 0, 1, ..., U_j^r$ be the number of observed T>C conversions. Let $h_j^r$ denote a hidden variable that indicates whether a read $j$ originates from a pre-existing RNA ($h_j^r = 0$), or a recently transcribed (i.e., newly synthesized) RNA ($h_j^r = 1$). Let $\rho^r \in [0, 1]$ be the $\frac{New}{Total}$ RNA ratio of region $r$.

Let $\ell \in [0,1]$ be the 4sU incorporation rate. Further, $\epsilon_+ \in [0,1]$ denotes the false-positive T>C sequencing error and $\epsilon_- \in [0,1]$ the false-negative C>X sequencing error. Let $\Theta = (\ell, \rho^r; r \in \mathcal{R})$ be the unknown parameters. Then,

$$P(o_j^r, h_j^r; \Theta) = P(h_j^r;\, \rho^r) \cdot P(o_j^r \mid h_j^r;\, U_j^r, \ell, \epsilon_+, \epsilon_-) \tag{19}$$

In Equation (19), $P(h_j^r; \rho^r) = Bernoulli(h_j^r; p = \rho^r)$ and

$$P(o_j^r \mid h_j^r; U_j^r, \ell, \epsilon_+, \epsilon_-) = \begin{cases} Bin(o_j^r;\, n = U_j^r, p = \epsilon_+) & \text{if } h_j^r = 0 \\ Bin(o_j^r;\, n = U_j^r, p = a) & \text{if } h_j^r = 1 \end{cases} \tag{20}$$

where $a = 1 - (\ell\epsilon_- + (1-\ell)(1-\epsilon_+))$ (Equation (8)). By taking logarithms, Equation (19) can be described as:

$$\log P(o_j^r, h_j^r; \Theta) = \begin{cases} \log(1-\rho^r) + \log\binom{U_j^r}{o_j^r} + o_j^r \log \epsilon_+ + (U_j^r - o_j^r)\log(1-\epsilon_+) & \text{if } h_j^r = 0 \\ \log \rho^r + \log\binom{U_j^r}{o_j^r} + o_j^r \log a + (U_j^r - o_j^r)\log(1-a) & \text{if } h_j^r = 1 \end{cases} \tag{21}$$

**E-step.**

Let $H = (h_j^r; j = 1, ..., J^r)$ and $H_{-1} = (h_j^r; j = 2, ..., J^r)$. Given a parameter set $\Theta' = (\rho^{r\prime}, \ell')$, the function $Q(\Theta, \Theta')$ has to be optimized with respect to $\Theta = (\rho^r, \ell)$:

$$\begin{aligned}
Q(\Theta;\Theta') &:= E_{P(H|O;\Theta')} \log P(O, H; \Theta) \\
&= \sum_H P(H \mid O; \Theta') \log P(O, H; \Theta) \\
&= \sum_{r\in\mathcal{R}} \sum_{H_{-1}^r} \sum_{h_1^r \in\{0,1\}} \left[P(H_{-1}^r \mid O_{-1}^r; \Theta') \cdot P(h_1^r \mid o_1^r; \Theta')\right] \\
&\quad \cdot \left[\log P(O_{-1}^r, H_{-1}^r; \Theta) + \log P(o_1^r, h_1^r; \Theta)\right] \\
&= \sum_{r\in\mathcal{R}} \sum_{h_1^r\in\{0,1\}} P(h_1^r \mid o_1^r; \Theta') \log P(o_1^r, h_1^r; \Theta) \\
&\quad + \sum_{r\in\mathcal{R}} \sum_{H_{-1}^r \in\{0,1\}^{J-1}} P(H_{-1}^r \mid O_{-1}^r; \Theta') \log P(O_{-1}^r, H_{-1}^r; \Theta) \\
&\overset{\text{induction}}{=} \sum_{r\in\mathcal{R}} \sum_{j=1}^{J^r} \sum_{h_j^r\in\{0,1\}} P(h_j^r \mid o_j^r; \Theta') \log P(o_j^r, h_j^r; \Theta)
\end{aligned} \tag{22}$$

Let

$$c_{j,h_j^r}^r := P(h_j^r \mid o_j^r; \Theta') = \frac{P(o_j^r \mid h_j^r; \ell') P(h_j^r; \rho'^r)}{\sum_{h_j^r\in\{0,1\}} P(o_j^r \mid h_j^r; \ell') P(h_j^r; \rho'^r)} \quad , \; j = 1, ..., J^r \tag{23}$$

where

$$\begin{aligned} C_0^r &= \sum_{j=1}^{J^r} c_{j,0}^r \\ C_1^r &= \sum_{j=1}^{J^r} c_{j,1}^r \end{aligned} \tag{24}$$

Further, if $U_j^r \geq 30$ (see Results section for determination of this threshold):

$$\begin{aligned} A &= \sum_{r\in\mathcal{R}} \sum_j c_{j,1}^r o_j^r \\ B &= \sum_{r\in\mathcal{R}} \sum_j c_{j,1}^r (U_j^r - o_j^r) \end{aligned} \tag{25}$$

Then, Equation (22) simplifies to:

$$\begin{aligned} Q(\Theta;\Theta') &= \sum_{r\in\mathcal{R}} \sum_j c_{j,0}^r \left[\log(1-\rho^r) + \log\binom{U_j^r}{o_j^r} + o_j^r \log\epsilon_+ + (U_j^r - o_j^r)\log(1-\epsilon_+)\right] \\ &+ \sum_{r\in\mathcal{R}} \sum_j c_{j,1}^r \left[\log\rho^r + \log\binom{U_j^r}{o_j^r} + o_j^r \log a + (U_j^r - o_j^r)\log(1-a)\right] \\ &= \sum_{r\in\mathcal{R}} C_0^r \log(1-\rho^r) + \sum_{r\in\mathcal{R}} C_1^r \log\rho^r + A\log a + B\log(1-a) + \text{const} \end{aligned} \tag{26}$$

**M-step.**

Taking the partial derivative of $Q$ in Equation (22) with respect to $\rho^r$ and equating this expression to zero yields:

$$\begin{aligned} 0 = \frac{\partial Q(\Theta;\Theta')}{\partial \rho^r} &= -\frac{C_0^r}{1-\rho^r} + \frac{C_1^r}{\rho^r} \\ \rho^r &= \frac{C_1^r}{C_0^r + C_1^r} = \frac{C_1^r}{J^r} \end{aligned} \tag{27}$$

Recall that $a = \ell(1-\epsilon_- - \epsilon_+) + \epsilon_+$ (Equation (8)), and take the partial derivative of $Q$ in Equation (22) with respect to $\ell$. Equating the resulting expression to zero yields:

$$\begin{aligned} 0 = \frac{\partial Q(\Theta;\Theta')}{\partial \ell} &= \frac{\partial Q(\Theta;\Theta')}{\partial a} \cdot \frac{\partial a}{\partial \ell} \\ &= \left(\frac{A}{a} - \frac{B}{1-a}\right) \cdot (1-\epsilon_- - \epsilon_+) \end{aligned} \tag{28}$$

Solving Equation (28) for $a$ (assuming $1-\epsilon_- - \epsilon_+ \neq 0$) and then solving for $\ell$ yields:

$$
\begin{aligned}
a &= \frac{A}{A+B} \\
\ell &= \frac{a - \epsilon_+}{1 - \epsilon_- - \epsilon_+} = \frac{(\frac{A}{A+B} - \epsilon_+)}{1 - \epsilon_- - \epsilon_+}
\end{aligned}
\tag{29}
$$

In addition, to stabilize the estimation, only regions with a certain coverage can be considered for the update step in Equation (25). In case of the SLAM-seq cell cycle data (Methods), a minimum of coverage of $J^r \geq 200$ reads per region was set.

### Variance-Stabilizing Transformation of $\frac{New}{Total}$ RNA Ratios

Halfpipe performs a variance-stabilization of the $\frac{New}{Total}$ RNA ratios $\rho$ as described in Müller *et al.* (2024). In brief, an arcsin transformation [25] is applied under the assumption that a 3'UTRs $\frac{New}{Total}$ RNA ratio $\rho^r$ and its corresponding total read count $J^r$ follow a binomial distribution given by $Bin(n = J^r, p = \rho^r)$. Subsequently, the arcsin-transformed $\frac{New}{Total}$ RNA ratios $arcsin\left(\sqrt{\rho^r}\right)$ can be approximated by a normal distribution with variance $\sigma^2 = \frac{1}{4 \cdot J^r}$. Here, $\sigma^2$ depends on the number of total reads $J^r$, which is accounted for during parameter fitting.

### Model Fitting Procedure

Equation (4)-(5) (two-compartment model) and Equation (6) (one-compartment model) are fitted using the arcsin-transformed $\frac{New}{Total}$ RNA ratios. First, fix a 3'UTR $r$. Let $q^r(t)$ be the arcsin-transformed $\frac{New}{Total}$ RNA ratio of the 3'UTR in measurement in either the nuclear or cytosolic compartment at time point $t$. Let $\hat{q^r}(t, \Theta)$ be the target $\frac{New}{Total}$ RNA ratio defined by $\Theta^r = (\tau^r + \nu^r, \lambda^r)$ (two-compartment model) or $\Theta^r = \delta^r$ (one-compartment model). The probability of observing the arcsin-transformed $\frac{New}{Total}$ RNA ratio $q^r(t)$ given the parameter set $\Theta$ can be formulated as:

$$
\begin{aligned}
P(q^r(t) \mid \Theta) &= \frac{1}{\sigma\sqrt{2\pi}} \cdot e^{-\frac{1}{2} \cdot \left(\frac{q^r(t) - \hat{q^r}(t,\Theta)}{\sigma}\right)^2} \\
&= \frac{\sqrt{4J^r}}{\sqrt{2\pi}} \cdot e^{-\frac{1}{2} \cdot 4J^r (q^r(t) - \hat{q^r}(t,\Theta))^2}
\end{aligned}
\tag{30}
$$

Subsequently, taking the logarithm results in:

$$
logP(q^r(t) \mid \Theta) = log\left(\frac{\sqrt{4J^r}}{\sqrt{2\pi}}\right) - 2J^r(q^r(t) - \hat{q^r}(t, \Theta))^2 \tag{31}
$$

Then, a suitable parameter set $\Theta^r$ is obtained by maximizing the sum of the log-likelihoods over all time points $T$ of a SLAM-seq time series $\sum_t^T logP(q^r(t) \mid \Theta^r)$ with respect to $\Theta^r$. In this case, $log\left(\frac{\sqrt{4J^r}}{\sqrt{2\pi}}\right)$ can be omitted from Equation (31) as it does not depend on $\Theta^r$.

The Markov chain Monte Carlo method (MCMC) Metropolis-Hastings [37] is used to sample $\Theta^r$. In case of the one-compartment model, a one-dimensional MCMC is generated. In case of the two-compartment model, $\tau^r + \nu^r$ and $\lambda^r$ are sampled jointly in a two-dimensional MCMC. A seed for the Markov chain is determined by global optimization (Differential Evolution algorithm [26]) to reduce burn-in time. The Markov chain samples 50,000 suitable parameter combinations. 95%-percentiles of these MCMC samples are defined as credibility intervals. Using median values of the MCMC samples, the final parameter set $\Theta^r$ is determined using local optimization (two-compartment model: Nelder-Mead algorithm [27]; one-compartment model: Brent algorithm [28]

### Figures

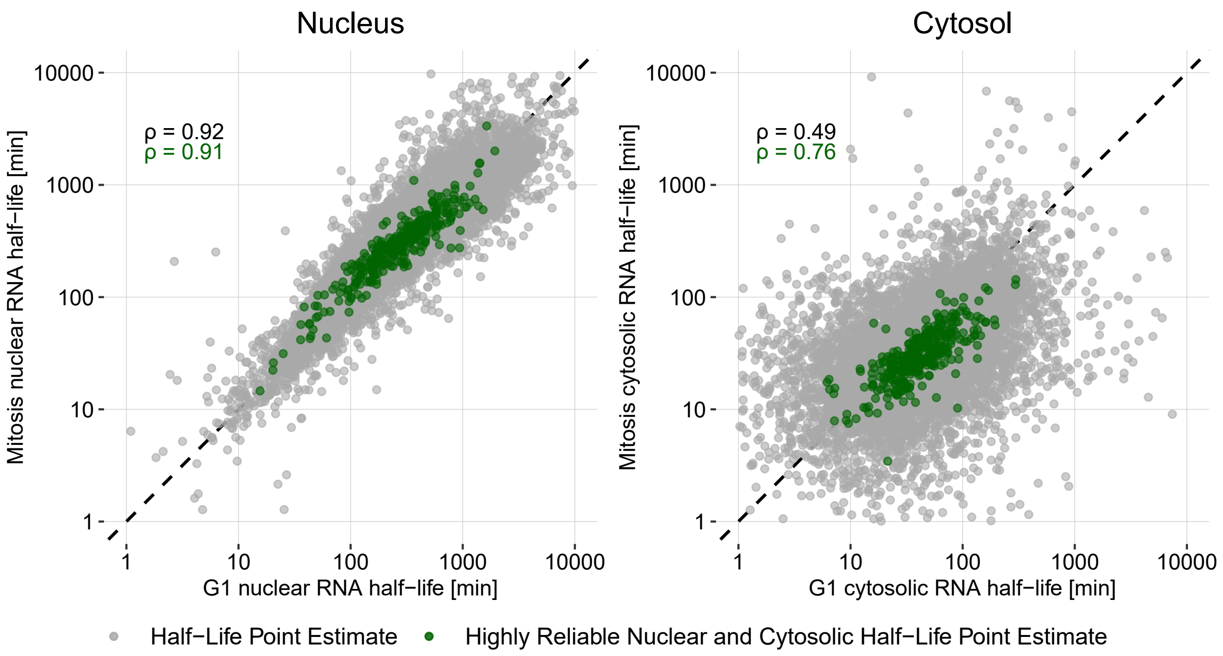


**Figure S1:** Scatterplot comparing compartment-specific RNA half-lives between G1 and Mitosis.
